# A natural fungal gene drive enacts killing through targeting DNA

**DOI:** 10.1101/2022.01.19.477016

**Authors:** Andrew S. Urquhart, Donald M. Gardiner

## Abstract

Fungal spore-killers are a class of selfish genetic elements that positively bias their own inheritance by killing non-inheriting gametes following meiosis. As killing takes place specifically within the developing fungal ascus, a tissue which is experimentally difficult to isolate, our understanding of the mechanisms underlying spore killers are limited. In particular, how these loci kill other spores within the fungal ascus is largely unknown. Here we overcome these experimental barriers by developing model systems in two evolutionary distant organisms, *Escherichia coli* (bacterium) and *Saccharomyces cerevisiae* (yeast). Using these systems, we show that the *Podospora anserina* spore killer protein Spok1 enacts killing through targeting DNA.

**Significance Statement:** Natural gene drives have shaped the genomes of many eukaryotes and recently have been considered for applications to control undesirable species. In fungi these loci are called spore-killers. Despite their importance in evolutionary processes and possible applications our understanding of how they enact killing is limited. We show that the spore killer protein Spok1, which has homologues throughout the fungal tree of life, acts via DNA disruption. Spok1 is only the second spore killer locus in which the cellular target of killing has been identified and is the first known to target DNA. We also show that the DNA disrupting activity of Spok1 is functional in both bacteria and yeast suggesting a highly conserved mode of action.

## Introduction

Gene drives are genetic elements that positively distort the frequency in which they are inherited. One group of meiotic gene drives are the evolutionarily diverse spore killer proteins found in ascomycete fungi. These gene drives distort normal mendelian inheritance by killing sibling progeny that do not inherit the locus [1]. Spore killers include het-s and Spok in *Podospora anserina* [2, 3]; Sk-1 and Sk-2 in *Neurospora* [4, 5], and *wtf* genes in *Schizosaccharomyces pombe* [6, 7].

Understanding how these gene drives function is important. Firstly, they likely influence genome evolution and structure, including mobile elements [8]. Secondly, uncovering their mechanisms might provide insight into conserved biological processes that they disrupt to enact killing [1]. And thirdly, they are potentially useful in pathogen control strategies. Gene drive approaches have already been widely investigated as control strategies for insect pests, notably malaria-transmitting mosquitoes [9]. The Spok1 spore killer from *Podospora anserina* has been shown to function heterologously in the dung-colonising fungus *Sordaria macrospora* [3] and the plant pathogen *Fusarium graminearum* [10]. The genetic control of plant-pathogenic fungi would decrease the need for the environmentally damaging fungicides currently used in agriculture.

Gene drive elements show considerable diversity. For example, the *Neurospora* Sk-2 and Sk-3 drives rely on two separate genes, encoding separate proteins to enact killing and resistance functions [11]. On the other hand, in some drive systems such as the *wtf* genes in *S. pombe* the killing and resistance functions are encoded by two separate proteins translated from alternative transcripts of the same gene [6, 7]. In the case of the Spok proteins in *P. anserina* only one transcript has been detected and it is not known how this single transcript can lead to both killing and resistance [12].

For the most part the cellular processes or structures targeted by the killing activity of spore killers is unknown. One exception is the *Het-s* locus (reviewed by [1]). Two alleles have been identified at this locus, *het-S* and *het-s*, encoding the proteins HET-S and HET-s respectively. Killing is enacted via interaction between these two proteins which induces a conformation change in the HET-S protein exposing a previously buried transmembrane domain. The altered HET-S protein perforates cells membranes killing the cell [13].

As with most other spore killers, the mechanism by which Spok proteins kill the developing gametes which did not inherit it is currently unknown. Bioinformatic analysis suggests that Spok proteins consists of three domains [12]. Experimental evidence suggests that the second domain, a putative nuclease, is required for killing activity and the third domain, a putative kinase is required for resistance. Specifically, a mutation (D667) within the third domain of Spok3 results in an allele which could not be transformed into *P. anserina*, suggesting that it was toxic even in vegetative tissue [12]. On the other hand, another mutation (K240) within the predicted catalytic core of the putative nuclease domain was found to abolish killing activity but not resistance [12]. Vogan *et al*. conjectured that a possible mode of action for the nuclease domain was the synthesis of a toxic diffusible metabolite [12].

A key difficulty in uncovering the mechanisms underlying spore killing is the limited availability of the relevant tissue (developing fungal asci). Indeed, in the case of het-S the mechanism was largely uncovered through examining its role in heterokaryon incompatibility, negating the need to examine developing asci. To overcome this issue, we sort to examine the activity of Spok1 in the experimentally amenable organism *Escherichia coli*. We here present evidence that the killing activity of Spok1^D680A^ protein (equivalent to Spok3^D667A^) is indeed active in *E. coli*, furthermore that this activity is mediated via the disruption of the chromosomal DNA and use the eukaryotic system of *Saccharomyces cerevisiae* to infer a mechanism involving DNA damage. Given that the killing activity is known to depend on a domain with similarity to restriction endonucleases, we suggest that direct modification (for example cleavage) of the genomic DNA by the nuclease domain represents a likely mechanism.

## Methods

### Strains used

#### *Escherichia coli* DH5α

F^−^ *endA1 glnV44 thi-1 recA1 relA1 gyrA96 deoR nupG purB20* φ80d*lacZ*ΔM15 Δ(*lacZYA-argF*)U169, hsdR17(*r*_*K*_ ^−^*m*_*K*_ ^+^), λ^−^. ***E. coli* Top10F’:** F’[*lacI*^q^Tn10(Tet^R^)] *mcrA* Δ(*mrr*-*hsdRMS-mcrBC*) φ80*lacZ*ΔM15 Δ*lacX74 recA1 araD139* Δ(*ara-leu*)*7697 galU galK rpsL endA1 nupG*. **NEB turbo** K-12 *glnV44 thi-1* Δ(*lac-proAB*) *galE15 galK16 R*(*zgb-210*::Tn*10*)Tet^S^ *endA1 fhuA2* Δ*(mcrB-hsdSM)5*(r_*K*_^−^ m_*K*_^−^) F′[*traD36 proAB*^+^ *lacI*^q^ *lacZΔM15*]. ***Saccharomyces cerevisiae* BY4742:** MATα *his3*Δ1 *leu2*Δ0 *lys2*Δ0 *ura3*Δ0.

### Molecular cloning

#### *E. coli* expression constructs

Plasmids were generated to express Spok genes (PLAUB44 Spok1^D680A^ or PLAUB51 Spok1^WT^) under the arabinose inducible P_BAD_ promoter [14]. The *araC*-P_BAD_ fragment was amplified from the genomic DNA of *E. coli* BL21(DE3) using primers AUB283 + AUB284. Two fragments of the yeast plasmid pYES2 backbone were amplified using primers AUB287 + DG1289; and primers DG1290 + AUB288.

Spok1^WT^ was amplified using primers AUB285 + AUB286 and Spok1^D680A^ was amplified in two fragments with AUB285 + AUB236 and AUB237 + AUB286. Oligonucleotide sequences are provided in Table S1. All PCR reactions were conducted using Q5 High-Fidelity 2X PCR Master Mix following the manufacturer’s directions (New England Biolabs). These fragments were transformed into *S. cerevisiae* by heat shock using a standard lithium acetate method [15] and were combined into a single plasmid via homologous recombination in the yeast cells [16]. The plasmid was rescued into chemically competent *E. coli* using the Zymoprep Yeast Plasmid Miniprep Kit (Zymo Research). Competent cells were prepared using the Inoue method [17].

#### *S. cerevisiae* expression construct

Spok1 coding DNA was amplified from PLAUB44 (Spok1^D680A^) or PLAUB51 (Spok1^WT^) [10] using AUB516 + AUB517 and cloned into the HindIII site of pYES2 using the NEBuilder HiFi DNA Assembly Master Mix (New England Biolabs) following the manufacturer’s directions. The primers were designed to include an optimal translation initiation site, including a synonymous C to T mutation in the second codon. This construct results in either Spok1^WT^ or Spok1^D680A^ expression under the galactose-inducible *GAL1* promoter of pYES2.

#### *E. coli* assays

*E. coli* containing the Spok1 plasmids were maintained on LB + carbenicillin + 0.2% glucose.

For induction of Spok1 expression the *E. coli* was first grown overnight in 5 ml LB + carbenicillin + glucose at 37°C with shaking. 50 μl of overnight culture was used to inoculate 50 ml of LB + carbenicillin in a 250 ml Erlenmeyer flask and allowed to grow for a further 4 h. At this point arabinose or glucose was added to 0.2% and incubation continued at 37°C either in the flask (for nucleic acid extraction) or diluted 1:1 with fresh LB media and incubated in a 96 well plate to acid monitor growth in a plate reader. Due to the faster growth rate of NEB Turbo cells this strain was diluted 1:10 rather than 1:1 with fresh LB. The plate reader assay was conducted in an EnVision Multimode Plate Reader with shaking (5 seconds, 900 rpm, 0.1 mm diameter, linear) every 10 min before reading OD at 595 nm.

To determine if Spok1^D680A^ expression merely arrested cells growth for effected permanent killing attempted to “rescue” the strains at various timepoints after arabinose induction. This was done by pipetting 10 μl of induced culture into 1 ml of LB + glucose then plating out onto LB + glucose + carbenicillin agar plates.

### RNA sequencing

*E. coli* strains containing plasmids PLAUB44 (Spok1^D680A^) or PLAUB51 (Spok1^WT^) were induced with arabinose as described above. RNA was extracted from 5 ml of *E. coli* culture at 45 min and 2 h 30 min, after induction for RNA sequencing. Four replicates for each construct at each timepoint were analysed (16 samples total). *E. coli* cultures were pelleted by centrifugation and then lyophilised before DNA was extracted using TRIzol reagent following the manufacturer’s instructions. The RNA was sequenced at the Australian Genome Research Facility. The libraries were prepared using the Illumina Stranded Total RNA Prep Ligation with Ribo-Zero Plus kit and sequenced on an Illumina NovaSeq 6000.

RNA sequence analysis was conducted using Galaxy [18]. Reads were mapped to the *E. coli* chromosome using STAR [19], reads mapping to each gene were counted using featureCount [20] and differentially expressed genes were determined using DEseq2 [21].

### DNA sequencing data and qPCR

20 ml of *E. coli* cultures were pelleted by centrifugation and then lyophilised before DNA was extracted using the Qiagen Plant Mini Kit. DNA was further purified by an ethanol precipitation step followed by resuspension in pure water. DNA quality was confirmed by gel electrophoresis and a nanodrop spectrophotometer. DNA was extracted at 45 min, 2 h 30 min, 4 h 30 min (3 biological replicates each) post arabinose induction for *E. coli* transformed with PLAUB44 (Spok1^D680A^) or PLAUB51 (Spok1^WT^).

The fC/ter ratio was determined using a hydrolysis probe based duplex qPCR assay designed to conform with the MIQE guidelines [22]. Primers TTCGATCACCCCTGCGTACA and CGCAACAGCATGGCGATAAC amplified part of the *gidA* gene located close to the origin [23]. This product was detected using a FAM labelled probe AUB458 6-FAM/ATGAGTGAT/ZEN/ATAACACGGCACCTGCTGG/IBFQ (IDT). Primers AUB484 + AUB485 were used to amplify part of the *dcp* gene located near the terminus. This product was detected using a Cy5 labelled probe Cy5/AACCCGCCC/TAO/TGCTGCTTATCGATAAC/IBRQ (IDT). qPCR reactions were conducted using Luna Universal Probe qPCR mastermix (New England Biolabs). Reactions were set up in a total volume of 10 ml containing 0.4 μM of each probe, 0.2 μM of each primer and approximately 100 ng of template DNA. Reactions were run on a Bio Rad CFX384 qPCR machine (initial denaturation of 95°C for 10 minutes followed by 95°C 10 sec, 58°C 10 sec, 72°C 15 sec for 40 cycles). The PCR efficacy was calculated using a 10-fold dilution series of DNA extracted from WT cells in late stationary phase (72 h). The oriC/ter of stationary phase cells is expected to be near 1 ([24-26]). *gidA* was found to amplify at 98.3% efficiency and *dcp* at 97.4% efficiency. Given the similar PCR efficiency between the two targets we did not adjust calculations to account for reaction efficiency.

The effect of Spok1^D680A^ expression on the *E. coli* chromosome was further explored via short-read whole-genome sequencing. DNA samples extracted from *E. coli* expressing Spok1^WT^ and Spok1^D680A^ 2 h 30 min after induction were sequenced on a NovaSeq 6000 generating 150 bp paired end reads at the Victorian Clinical Genetics Services. The resultant reads were mapped to the *E. coli* chromosome using Bowtie 2 [27] in Galaxy, coverage across the chromosome was calculated from the resultant BAM file using bamCoverage [28] using 1 kb windows.

### *RAD51* deletion in *S. cerevisiae*

The *LEU2* selectable marker was amplified from plasmid pGAD-c1 [29] using primers AUB528 + AUB529. The resultant PCR product was used to delete the *RAD51* gene via homologous recombination in *S. cerevisiae* strain BY4742. Transformation was conducted using LiAc/PEG [15] and deletion of the gene was confirmed using primers AUB488 + AUB489, which amplify across the mutated region.

### *S. cerevisiae* growth assay

To determine the effect of Spok1^D680A^ expression in *S. cerevisiae* a spot assay on agar plates was employed. Strains carrying PLAUB90 (Spok1^WT^) or PLAUB91 (Spok1^d680A^) were streaked on SD without uracil with 2% glucose or galactose and grown for 48 h at 30°C. Colonies were picked into sterile water, serially diluted, and plated out onto SD without uracil with 2% glucose or galactose plates.

### *RNR3* promoter GFP reporter strain

GFP was introduced into the *S. cerevisiae RNR3* locus. GFP was amplified from plasmid PLAU17 [30] with primers AUB536 + AUB537 the *LEU2* gene was amplified with primers AUB538 + AUB539 from plasmid pGAD-c1 [29]. Both products were simultaneously transformed into *S. cerevisiae* strain BY4742. Transformants were screened for hydroxyurea (HU) inducible GFP expression (SC with 50 mM HU versus no HU). GFP expression was quantified using a Biotek Cytation 1 plate imager at 12 h hours post induction on a microscope slide. The fluorescence of individual cells was quantified using Image J software. Cell were pelleted via centrifugation and resuspended in water before examination to minimise background fluorescence.

The Spok1 plasmids were introduced into the RNR3-GFP strain and GFP fluorescence was measured following 12 h growth in SD without uracil with galactose.

## Results

### A Spok1 allele carrying a mutated resistance domain (Spok1^D680A^) is toxic to *E. coli*

Inspired by the autoactive version of Spok3 being functional in vegetative cells of *P. anserina* [12] we reasoned that killing may target a conserved biological process and sought to identify a highly tractable system in which toxicity could be assessed. Such a system requires tightly regulable gene expression, which in *E. coli* can be achieved with the arabinose-inducible and glucose-repressible P_BAD_ promoter [14]. A protein alignment revealed that the Spok3 residue D667 previously found to be essentially for resistance activity [12] corresponds to residue D680 in Spok1 (Figure 1A) and we created the equivalent mutated allele in Spok1 (hereafter termed Spok1^D680A^). When cloned under the control of the P_BAD_ promoter, the wildtype and autoactive version of Spok1 grew similarly under repressive (high glucose) conditions. Expression of Spok1^D680A^ caused *E. coli* growth to stop approximately 100 minutes after arabinose induction (Figure 1). In contrast cells expressing Spok1^WT^ continue to grow. This effect was observed in two commonly used *E. coli* strains Top10F’ (arabinose non utilising) and DH5α (arabinose utilising).

**Figure 1:**
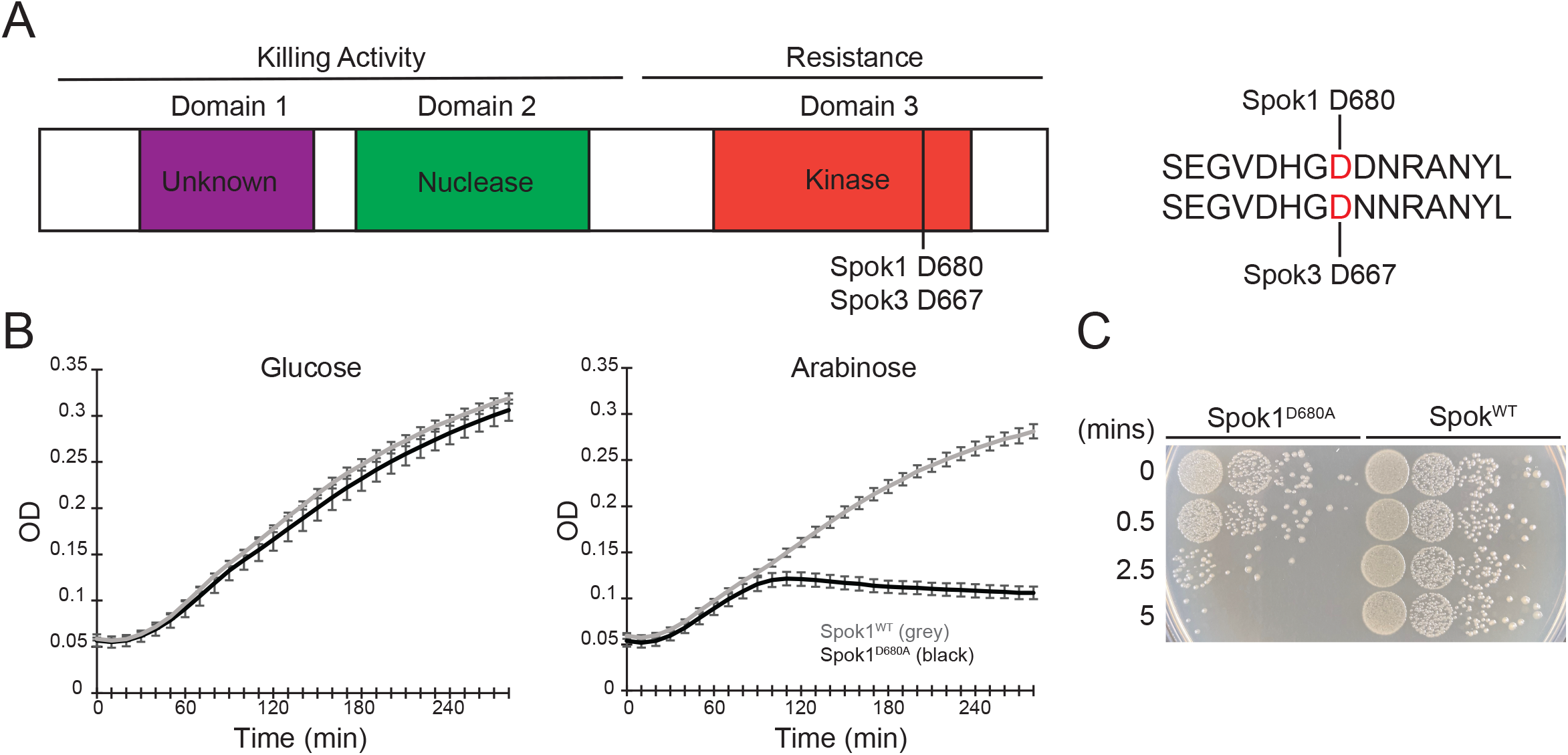
**A)** Representation of the domain structure of Spok proteins, according to [12]. The spore killing activity of the Spok proteins requires the nuclease (second) and possibly first domains. Host resistance is mediated by the third domain, a putative kinase. Experimental evidence suggests that mutation of aspartic acid (D) 667 to alanine (A) in Spok3 results is a toxic allele possessing only killing activity [12]. Protein alignments showed that this corresponds to residue D680 in Spok1. **B)** Growth curve of *E. coli* expressing Spok1^WT^ and Spok1^D680A^ after induction by arabinose. Error bars represent ± 1 standard deviation. **C)** 10-fold dilution series of induced cells at various timepoints after induction passaged back onto high glucose concentrations and incubated for 24 hours to allow colonies to develop.

We next sought to determine if the expression of Spok1^D680A^ was merely an inhibition of growth or indeed genuine killing. To this end we attempted to rescue arabinose induced cells by rapidly diluting them into high concentrations of glucose to supress gene expression. This assay showed that the *E. coli* cells expressing Spok1^D680A^ quickly lose viability after induction with an almost complete loss of viability after just 5 minutes in inducive conditions (Figure 1C).

### DNA metabolism genes are differentially regulated in response to Spok1^D680A^ expression

To determine the mechanisms underlying the killing in *E. coli* we conducted RNA sequencing on induced *E. coli* expressing either Spok1 or Spok1^D680A^. Timepoints for RNA sequencing were chosen based on the OD curves rather than the rescue assay as OD readings are likely to better reflect the timing of physiological changes in the cell, even though the Spok1^D680A^-expressing cells were committed to death earlier. At an early timepoint (45 min), expression changes were relatively minor compared to the later timepoint (2 h 30 min) (Figure 2A). Only 47 genes showed statistically significant (at p<0.05) and greater than 2-fold change at 45 min compared to 1076 genes at 2 h 30 min (Table S2).

**Figure 2:**
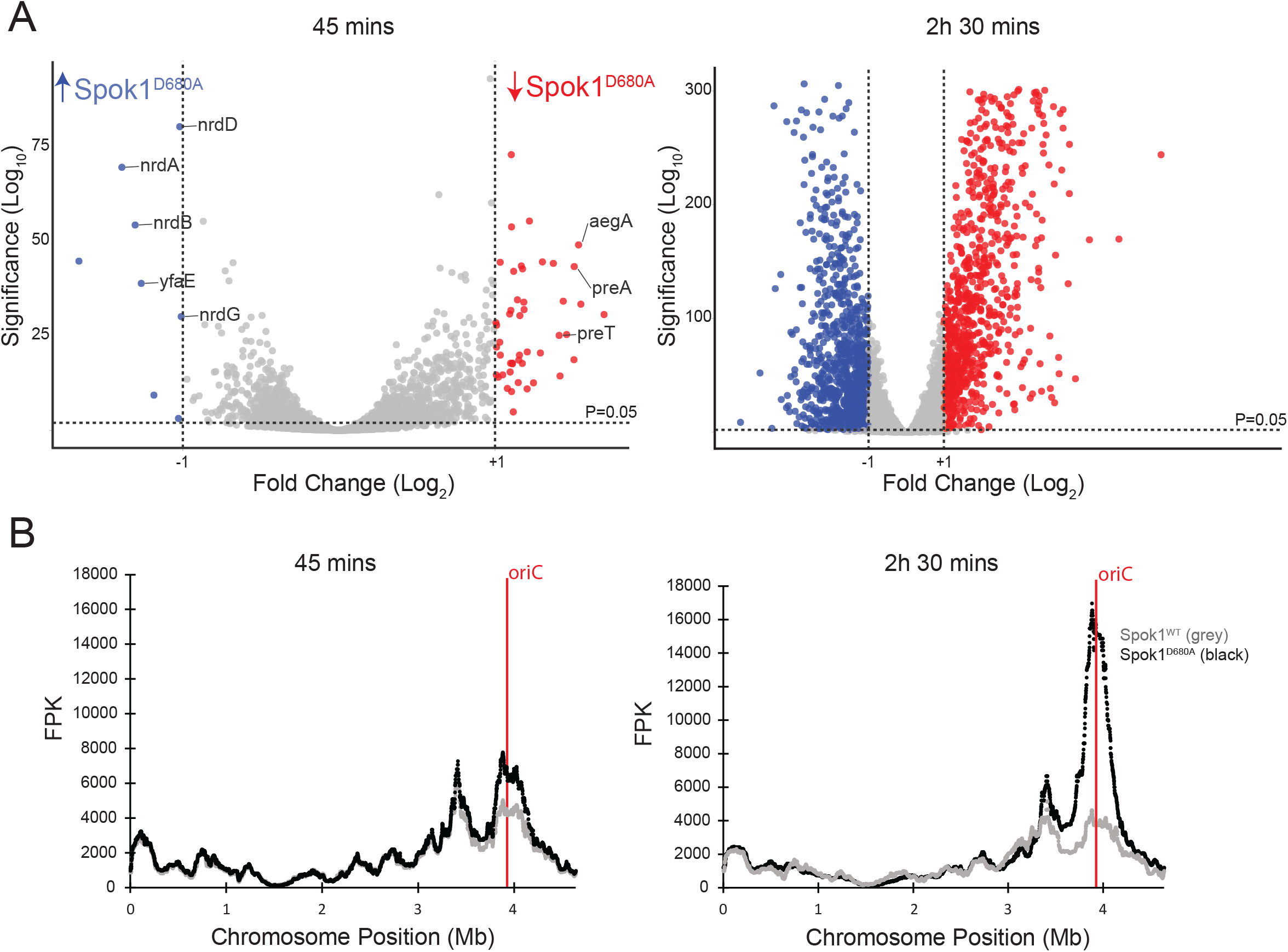
**A)** Volcano plots showing RNA seq data at 45 minutes and 2 hours 30 minutes. Blue genes are more highly expressed in *E. coli* expressing Spok1^D680A^ and red genes are more highly expressed (greater than 2-fold change) in *E. coli* expressing Spok1^WT^. Specific genes involved in DNA metabolism that are differentially regulated at 45 min post induction are annotated. **B)** Median gene expression (100 gene windows) along the *E. coli* chromosome. Increased expression in genes proximal to the oriC in *E. coli* expressing Spok1^D680A^ (compared to cells expressing Spok1^WT^) is observed at both 45 minutes and 2 hours 30 minutes but is more pronounced at the latter timepoint.

Examination of the 47 genes showing altered regulation at 45 min revealed at least 8 genes involved in nucleotide metabolism including *nrdA, nrdB, nrdD, nrdG, yfaE, aegA, preA* and *preT*. Four of these genes (which were all upregulated in the Spok1^D680A^ expressing strain) encode subunits of the two *E. coli* ribonucleotide reductases (RNR). RNRs convert ribonucleotides to deoxyribonucleotides which are essential for DNA synthesis. NrdA and NrdB form the aerobic RNR in *E. coli* and the neighbouring gene *yfaE* may play a role in the functioning of this enzyme [31]. NrdD and NrdG form the anaerobic RNR [32]. The down-regulated genes included genes required for the degradation of pyrimidines and purines. Namely PreA and PreT catalyse the reduction of uracil to 5,6-dihydrouracil in the breakdown of pyrimidine bases [33]. AegA is involved in the breakdown of purine nucleotides through the degradation of urate [34].

By the 2 h 30 min timepoint the extent of gene expression changes rendered the identification of individual genes impractical (Figure 2A). However, mapping gene expression along the chromosome revealed an upregulation in genes surrounding the origin of replication in the strain expressing Spok^D680A^ relative to the strain expressing Spok1^WT^ (Figure 2B). This difference was present at 45 min but more pronounced at 2 h 30 min.

### Spok1^D680A^ disrupts chromosome replication resulting in increased copy number proximal to the origin

DNA replication in *E. coli* proceeds bidirectionally from the origin of replication (oriC) to the terminus (ter) and new replication forks are initiated before previous rounds of DNA replication have completed, meaning that in actively growing cells there will be more copies of the DNA closer to the initiation of the DNA replication fork (Figure 3A) [35]. We hypothesised that apparent up-regulation of gene expression surrounding the origin of replication was a result of a corresponding increased DNA copy number in a gradient from the oriC to ter loci. We therefore determined the ratio of cellular DNA between the oriC and ter loci using quantitative PCR (Figure 3B). This revealed that induction of Spok1^D680A^ resulted in an oriC/ter ratio of approximately 50:1 compared to approximately 2:1 in the strain expressing Spok1^WT^. The increase in DNA copy number surrounding the origin was confirmed by whole genome DNA sequencing (at 2 h 30 min) as this showed increased relative copy number in a region of approximately 1 million bases peaking at the origin of replication (Figure 3C).

**Figure 3:**
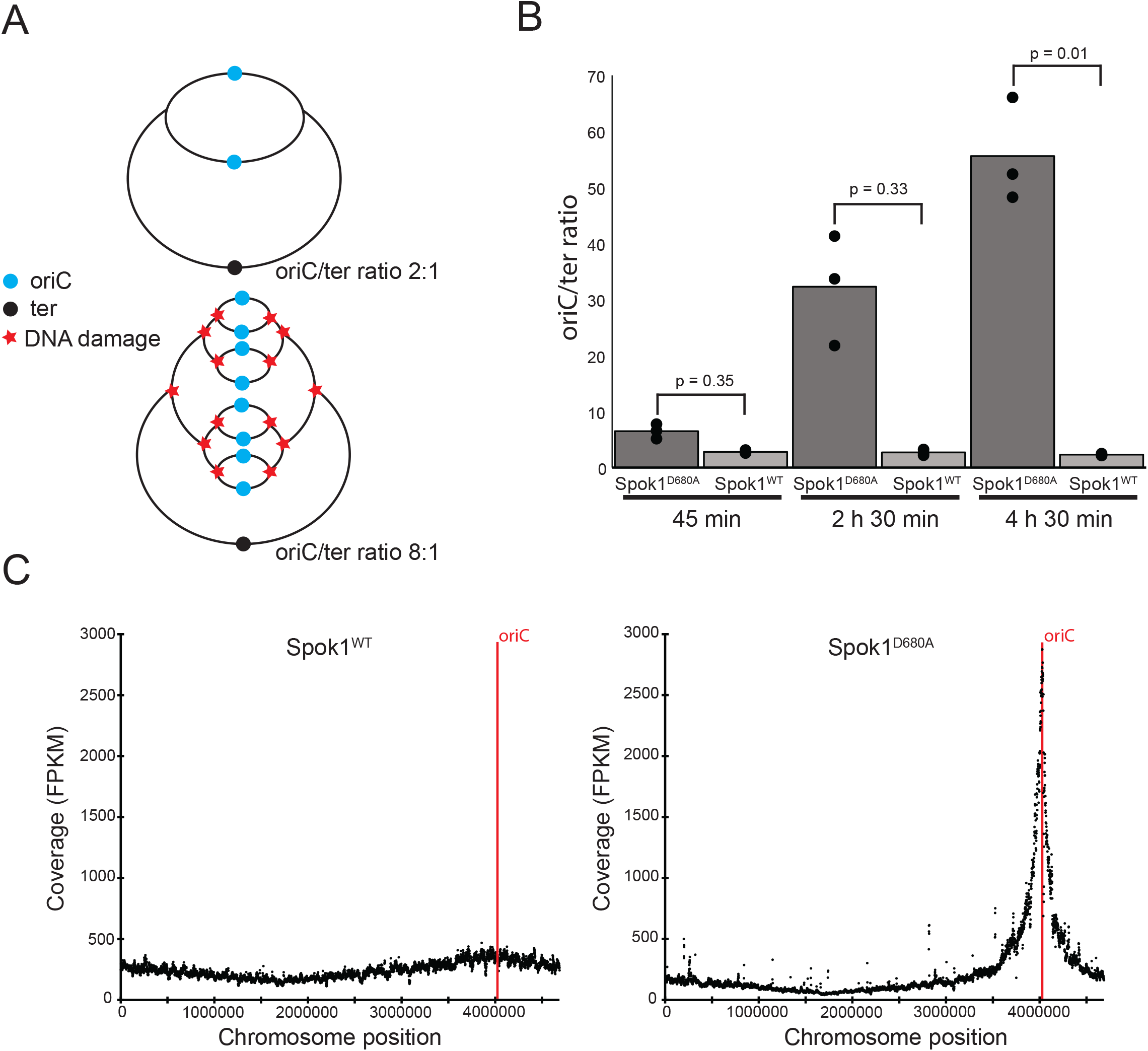
**A)** Diagram displaying the effect of DNA damage on oriC/ter ratio. If DNA damage prevents the replication form progressing fewer replication forks will reach the terminus of the chromosome. As replication forks continue to be initiated but fail to complete replication the ratio between oriC and ter DNA will increase. **B)** qPCR analysis of the oriC/ter ratio following induction of either Spok1^WT^ or Spok1^D680A^. DNA sequencing read depth compared to chromosomal position at 2 h 30 min following induction of either Spok1^WT^ or Spok1^D680A^. **C)** Read depth (FPKM) of Illumina DNA sequencing reads mapped to the *E. coli* chromosome, each point represents a 1kb genomic window.

### Spok1 also targets DNA to enact killing activity in a eukaryote

While *E. coli* is a tractable system for analysing the mechanisms of the Spok proteins, there are fundamental differences in the cell biology between the fungi in which Spok genes are found and prokaryotes, not least the enveloped nucleus in eukaryotes in which the Spok proteins are thought to reside [3]. To understand if our identification of DNA as the target of Spok proteins also occurred in eukaryotes we utilised baker’s yeast as it lacks endogenous copies of Spok genes but shares many fundamental aspects of eukaryote biology with filamentous ascomycetes. Indeed expression of Spok^D680A^ in *S. cerevisiae* reduced growth rate (Fig 4C).

**Figure 4:**
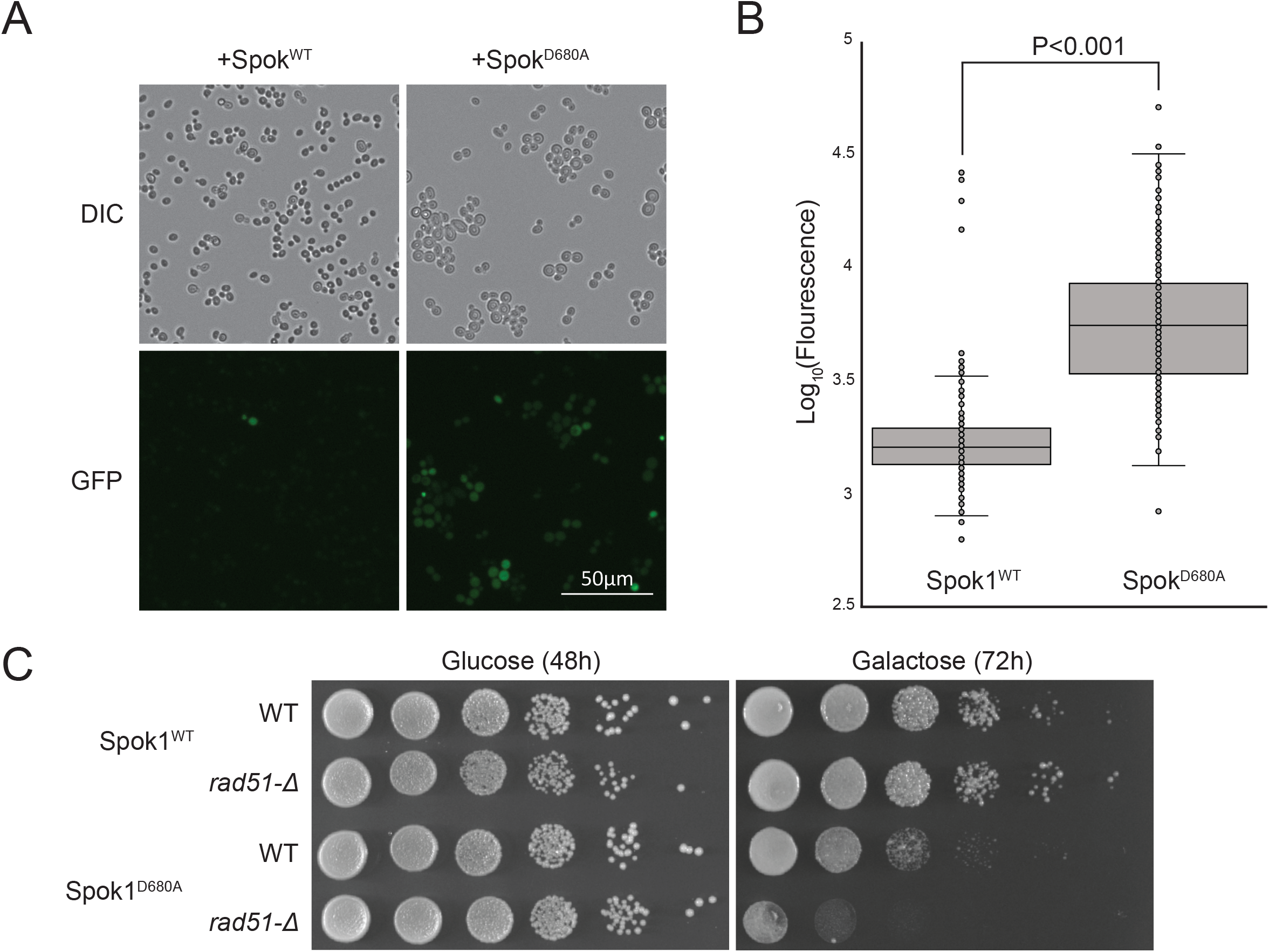
**A and B)** Expression of Spok1^D680A^ in a DNA damage responsive RNR3:GFP reporter strain induced expression of GFP. Data points represent fluorescence intensity of induvial cells (n= 256). **C)** Spot assay of *S. cerevisiae* WT and *rad51*Δ expressing Spok^WT^ or Spok1^D680A^ under a galactose inducible promoter. Cells were passage from glucose media onto either glucose or galactose media and the amount of growth following incubation at 30°C was observed. The *rad51*Δ mutant was hypersensitive to Spok1^D680A^ expression.

To determine if DNA was a target of Spok1 we sought to assay whether Spok1^D680A^ was triggering DNA damage in yeast. To do this we made developed an RNR3:GFP strain to monitor expression of *RNR3* which is known to be responsive to DNA damage such as hydroxyurea as previously reported (data not shown, [36]). Induction of Spok1^D680A^ in this reporter strain (Fig 4A and B) demonstrates upregulation of *RNR3* compared to expression of Spok1^WT^ which presumably is in response to DNA damage.

Having established that DNA damage was most likely also occurring when Spok1^D680A^ was expressed, we hypothesised that mutants impaired in their ability to repair DNA damage would show hypersensitivity to Spok1^D680A^ expression. Rad51 deletion mutants show increased sensitivity to DNA damaging agents with Rad51 acting as a DNA recombinase in pathways for repairing DNA damage [37]. As expected, the *rad51*Δ mutant to showed hypersensitivity to Spok^D680A^ expression (Fig 4C).

**Figure S1.**
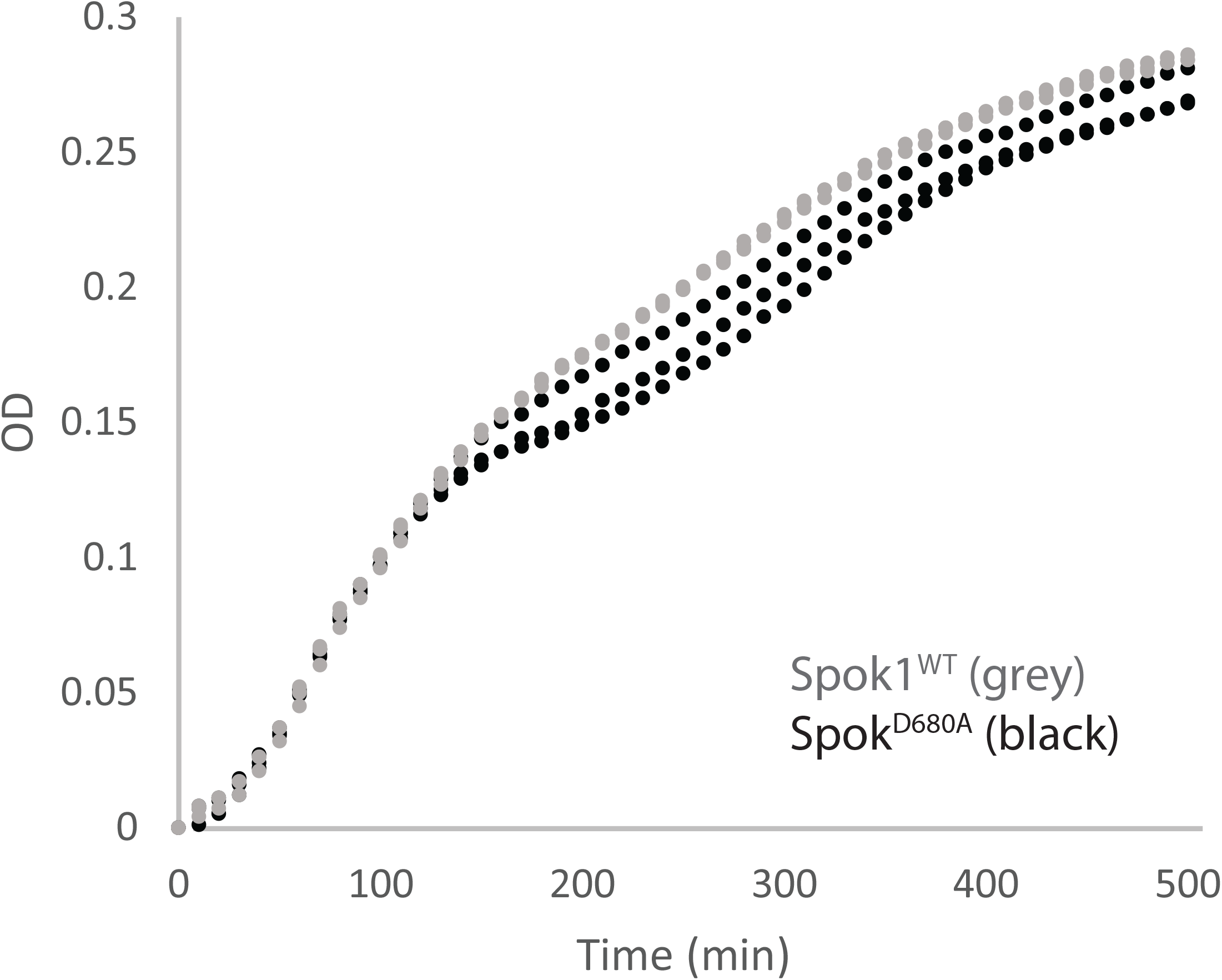
Expression of Spok1^WT^ in an *E. coli* RecA+ strain (NEB Turbo) resulted in less inhibition that in recA-strains (compare to Figure 1B in which growth ceased at approximately 100 minutes following induction of Spok1^D680A^).

## Discussion

The mechanisms underlying the gene drive mechanisms of fungal spore killers are poorly understood. We here provide several lines of evidence that the Spok1 protein of *Podospora* acts through DNA damage. Given the complexity of the relevant tissue (developing fungal ascus in the context of structural components of the perithecia) we employed heterologous model systems. We demonstrated that a mutated version of the Spok1 protein lacking resistance function (Spok1^D680A^) was toxic in the model organisms *E. coli* and *S. cerevisiae* (Figures 1B and 4A). These systems provide obvious experimental advantages over examining meiotic drive *in situ* during fungal crosses. Because the protein was functional in organisms as evolutionary distant as *E. coli* and yeast we hypothesised that Spok killing activity must target a highly conserved structure/pathway. Furthermore, the conservation of key residues between Spok1 and other Spok proteins, presumably the mode of action described here will be conserved in this gene family.

Firstly, we took a mechanism-neutral approach in *E. coli* and conducted RNA-sequencing on *E. coli* cultures induced to express Spok1^D680A^. At an early timepoint (45 min), several genes involved in DNA metabolism were differentially regulated (Figure 2A). These included several RNR subunits. *E. coli* RNR genes are known to be regulated in response to DNA damage [38]. Of note is that the RNA sequencing analysis was conducted in a *recA* mutant for improved plasmid stability. RecA is an important regulator of DNA repair pathways so many DNA-damage inducible genes will be unresponsive in this strain [39]. The altered regulation of genes involved in DNA metabolism at 45 minutes post-induction provided initial evidence that DNA metabolism was the target of Spok1 killing activity.

Examination of a later timepoint (2 h 30 min post induction) showed an unexpectedly strong expression of genes within a region of ∼1Mbp centred around the origin of replication. Gene expression levels are known to correlate with chromosomal position in *E. coli* with highest expression closest to the origin [40]. However, in *E. coli* expressing Spok1^D680A^ this effect was amplified (Figure 2B).

With evidence for specific DNA-metabolism related genes being regulated at the 45 min timepoint, we hypothesised that this effect might be due to an underlying perturbation in corresponding DNA copy number. The circular *E. coli* chromosome replicates bidirectionally from the origin to the terminator. Because new replication forks are initiated before previous replication forks have completed this results in a ratio of oriC/ter greater than one in actively dividing cells [25].

Treatments which block the progression of the replicon e.g., hydroxyurea (which inhibits RNR and damages DNA) or antibiotics such as trimethoprim (which inhibit DNA synthesis) are known to increase the oriC/ter ratio [41, 42]. qPCR and whole-genome Illumina sequencing confirmed that the RNA-seq expression patterns were a result of DNA perturbations (Figure 3).

Given the evolutionary distance between bacteria and the eukaryotes in which Spok proteins occur natively, we decided to induce expression of Spok1^D680A^ in the yeast *S. cerevisiae*. As in *E. coli*, expression of Spok1^D680A^ protein proved toxic to the yeast cells. We took two difference approaches to demonstrate that the killing of *S. cerevisiae* occurred through genotoxic stress. The first was to express Spok1^D680A^ in an *S. cerevisiae* RNR3:GFP reporter strain. Rnr3 is a ribonucleotide reductase gene that is expressed in response to DNA damage [36]. Induction of Spok1^D680A^ induced GFP fluorescence in this strain (Figure 4A and B). Secondly, we expressed Spok1^D680A^ in a *rad51*Δ mutant. This mutant is known to be deficient in DNA repair and was hypersensitive to Spok1^D680A^ (Figure 4C). Conversely, a DNA repair proficient (*recA+*) *E. coli* strain proved more resistant to killing than the *recA-*cloning strains used elsewhere in this study (Supp Fig 1).

A limitation of this study is that we have not examined the activity of Spok1 *in situ* during fungal crosses to demonstrate the occurrence of DNA damage within a fungal ascus, as such an experiment is technically challenging. The D680A mutation behaves in the same way in both *E. coli* and *S. cerevisiae*, with evidence of DNA stress occurring in both species, suggests that the same mode of action will occur in the fungal ascus. The common response to Spok1^D680A^ expression in these two distantly related species is consistent with a highly conserved molecular target such as genomic DNA.

The exact mechanism underlying the effect of Spok1^D680A^ on DNA is still unclear. The killing activity of Spok1 proteins has previously been shown to require a catalytically active nuclease domain with similarity to type I restriction endonucleases [12]. Also consistent with DNA targeting, Grognet et al. showed that Spok proteins localise to the fungal nucleus in developing asci [3]. We thus suggest that the simplest mechanism for Spok1 killing, consistent with our data, the nuclear localisation and the domain content of the protein, would be cleavage (or other direct damage) of the chromosomal DNA by the nuclease domain of Spok1. Other toxins are known to work via enzymatic DNA damage for example Cytolethal distending toxin (CDT) produced by certain bacterial pathogens causes double-stranded DNA breaks in mammalian cells [43].

There are parallels to bacterial restriction modification (RM) systems which are known to behave as selfish elements [44]. They consist of a restriction enzyme and a methyltransferase that methylates the target site and thus protects the genome. However, if the selfish RM element is lost from the cell (e.g. if carried on an unstable plasmid) then the cell is killed (reviewed by [45]). This is curiously reminiscent of fungal gene drives in which spores not inheriting the drive element are killed.

However, in the case of Spok proteins the resistance activity is encoded by a putative kinase domain of the same protein, not a separate methyltransferase enzyme, so the mechanism underlying resistance remains unclear. We envisage that the *E. coli* and *S. cerevisiae* models reported here will enable further studies into the activity of Spok proteins including how resistance is mediated.

## Supporting information

Table S2 DEseq2 Output

## Data availability

RNA and DNA sequencing reads are available under BioProject PRJNA798172.

## Acknowledgements

ASU was supported by a CSIRO Early Research Career Postdoctoral Fellowship. DMG was supported by the Australian Research Council Research Hub for Sustainable Crop Protection (project number IH190100022) and by the Australian Government.

---

**Table S1:**
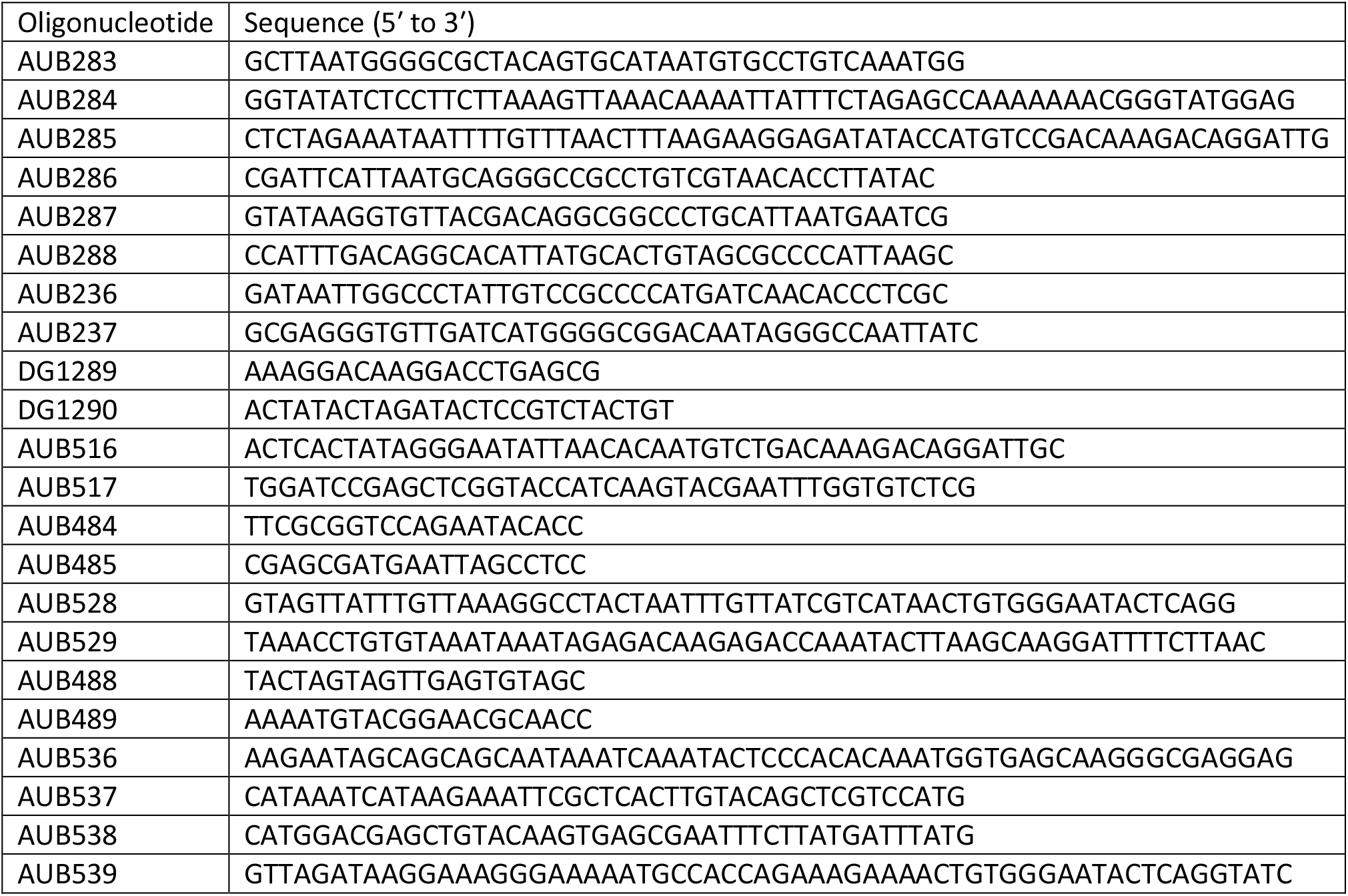
Oligonucleotides designed in this study:

## References

1. Zanders, S., and Johannesson, H. (2021). Molecular mechanisms and evolutionary consequences of spore killers in ascomycetes. Microbiol Mol Biol Rev 85, e0001621.

2. Dalstra, H.J.P., Swart, K., Debets, A.J.M., Saupe, S.J., and Hoekstra, R.F. (2003). Sexual transmission of the [Het-s] prion leads to meiotic drive in Podospora anserina. Proceedings of the National Academy of Sciences 100, 6616–6621.

3. Grognet, P., Lalucque, H., Malagnac, F., and Silar, P. (2014). Genes that bias mendelian segregation. PLoS genetics 10, e1004387.

4. Svedberg, J., Vogan, A.A., Rhoades, N.A., Sarmarajeewa, D., Jacobson, D.J., Lascoux, M., Hammond, T.M., and Johannesson, H. (2021). An introgressed gene causes meiotic drive in Neurospora sitophila. Proceedings of the National Academy of Sciences 118, e2026605118.

5. Rhoades, N.A., Harvey, A.M., Samarajeewa, D.A., Svedberg, J., Yusifov, A., Abusharekh, A., Manitchotpisit, P., Brown, D.W., Sharp, K.J., Rehard, D.G., et al. (2019). Identification of rfk-1, a meiotic driver undergoing RNA editing in Neurospora. Genetics 212, 93–110.

6. Hu, W., Jiang, Z.D., Suo, F., Zheng, J.X., He, W.Z., and Du, L.L. (2017). A large gene family in fission yeast encodes spore killers that subvert Mendel’s law. eLife 6.

7. Nuckolls, N.L., Bravo Núñez, M.A., Eickbush, M.T., Young, J.M., Lange, J.J., Yu, J.S., Smith, G.R., Jaspersen, S.L., Malik, H.S., and Zanders, S.E. (2017). wtf genes are prolific dual poison-antidote meiotic drivers. eLife 6.

8. Vogan, A.A., Ament-Velásquez, S.L., Bastiaans, E., Wallerman, O., Saupe, S.J., Suh, A., and Johannesson, H. (2021). The Enterprise, a massive transposon carrying Spok meiotic drive genes. Genome Res. 31, 789–798.

9. Nolan, T. (2021). Control of malaria-transmitting mosquitoes using gene drives. Philosophical Transactions of the Royal Society B: Biological Sciences 376, 20190803.

10. Gardiner, D.M., Rusu, A., Barrett, L., Hunter, G.C., and Kazan, K. (2020). Can natural gene drives be part of future fungal pathogen control strategies in plants? New Phytologist 228, 1431–1439.

11. Hammond, T.M., Rehard, D.G., Xiao, H., and Shiu, P.K.T. (2012). Molecular dissection of Neurospora spore killer meiotic drive elements. Proceedings of the National Academy of Sciences 109, 12093–12098.

12. Vogan, A.A., Ament-Velásquez, S.L., Granger-Farbos, A., Svedberg, J., Bastiaans, E., Debets, A.J., Coustou, V., Yvanne, H., Clavé, C., Saupe, S.J., et al. (2019). Combinations of Spok genes create multiple meiotic drivers in Podospora. eLife 8.

13. Seuring, C., Greenwald, J., Wasmer, C., Wepf, R., Saupe, S.J., Meier, B.H., and Riek, R. (2012). The mechanism of toxicity in HET-S/HET-s prion incompatibility. PLoS Biology 10, e1001451.

14. Guzman, L.M., Belin, D., Carson, M.J., and Beckwith, J. (1995). Tight regulation, modulation, and high-level expression by vectors containing the arabinose P<sub>BAD </sub>promoter. Journal of Bacteriology 177, 4121–4130.

15. Gietz, R.D., and Schiestl, R.H. (2007). High-efficiency yeast transformation using the LiAc/SS carrier DNA/PEG method. Nat. Protoc. 2, 31–34.

16. Oldenburg, K. (1997). Recombination-mediated PCR-directed plasmid construction in vivo in yeast. Nucleic Acids Research 25, 451–452.

17. Inoue, H., Nojima, H., and Okayama, H. (1990). High efficiency transformation of Escherichia coli with plasmids. Gene 96.

18. Jalili, V., Afgan, E., Gu, Q., Clements, D., Blankenberg, D., Goecks, J., Taylor, J., and Nekrutenko, A. (2020). The Galaxy platform for accessible, reproducible and collaborative biomedical analyses: 2020 update. Nucleic Acids Research 48, W395–W402.

19. Dobin, A., Davis, C.A., Schlesinger, F., Drenkow, J., Zaleski, C., Jha, S., Batut, P., Chaisson, M., and Gingeras, T.R. (2013). STAR: ultrafast universal RNA-seq aligner. Bioinformatics 29, 15–21.

20. Liao, Y., Smyth, G.K., and Shi, W. (2014). featureCounts: an efficient general purpose program for assigning sequence reads to genomic features. Bioinformatics 30, 923–930.

21. Love, M.I., Huber, W., and Anders, S. (2014). Moderated estimation of fold change and dispersion for RNA-seq data with DESeq2. Genome biology 15.

22. Bustin, S.A., Benes, V., Garson, J.A., Hellemans, J., Huggett, J., Kubista, M., Mueller, R., Nolan, T., Pfaffl, M.W., Shipley, G.L., et al. (2009). The MIQE guidelines: Minimum information for publication of quantitative real-time PCR experiments. Clinical Chemistry 55, 611–622.

23. Riber, L. (2006). Hda-mediated inactivation of the DnaA protein and dnaA gene autoregulation act in concert to ensure homeostatic maintenance of the Escherichia coli chromosome. Genes & Development 20, 2121–2134.

24. Brochu, J., Vlachos-Breton, É., Sutherland, S., Martel, M., and Drolet, M. (2018). Topoisomerases I and III inhibit R-loop formation to prevent unregulated replication in the chromosomal Ter region of Escherichia coli. PLoS genetics 14, e1007668.

25. Haugan, M.S., Charbon, G., Frimodt-Møller, N., and Løbner-Olesen, A. (2018). Chromosome replication as a measure of bacterial growth rate during Escherichia coli infection in the mouse peritonitis model. Scientific Reports 8.

26. Simmons, L.A., Breier, A.M., Cozzarelli, N.R., and Kaguni, J.M. (2004). Hyperinitiation of DNA replication in Escherichia coli leads to replication fork collapse and inviability. Molecular Microbiology 51, 349–358.

27. Langmead, B., and Salzberg, S.L. (2012). Fast gapped-read alignment with Bowtie 2. Nature Methods 9, 357–359.

28. Ramírez, F., Ryan, D.P., Grüning, B., Bhardwaj, V., Kilpert, F., Richter, A.S., Heyne, S., Dündar, F., and Manke, T. (2016). deepTools2: a next generation web server for deep-sequencing data analysis. Nucleic Acids Research 44, W160–W165.

29. James, P., Halladay, J., and Craig, E.A. (1996). Genomic libraries and a host strain designed for highly efficient two-hybrid selection in yeast. Genetics 144, 1425–1436.

30. Idnurm, A., Urquhart, A.S., Vummadi, D.R., Chang, S., Van de Wouw, A.P., and López-Ruiz, F.J. (2017). Spontaneous and CRISPR/Cas9-induced mutation of the osmosensor histidine kinase of the canola pathogen Leptosphaeria maculans. Fungal Biol. Biotechnol. 4, 12.

31. Wu, C.H., Jiang, W., Krebs, C., and Stubbe, J. (2007). YfaE, a ferredoxin involved in diferric-tyrosyl radical maintenance in Escherichia coli ribonucleotide reductase. Biochemistry 46, 11577–11588.

32. Garriga, X., Eliasson, R., Torrents, E., Jordan, A., Barbé, J., Gibert, I., and Reichard, P. (1996). nrdD and nrdG genes are essential for strict anaerobic growth of Escherichia coli. Biochemical and Biophysical Research Communications 229, 189–192.

33. Hidese, R., Mihara, H., Kurihara, T., and Esaki, N. (2011). Escherichia coli dihydropyrimidine dehydrogenase Is a novel NAD-dependent heterotetramer essential for the production of 5,6-dihydrouracil. Journal of Bacteriology 193, 989–993.

34. Iwadate, Y., and Kato, J.I. (2019). Identification of a formate-dependent uric acid degradation pathway in Escherichia coli.. Journal of Bacteriology 201, e00573–00518.

35. Wang, J.D., and Levin, P.A. (2009). Metabolism, cell growth and the bacterial cell cycle. Nature Reviews Microbiology 7, 822–827.

36. Suzuki, H., Sakabe, T., Hirose, Y., and Eki, T. (2017). Development and evaluation of yeast-based GFP and luciferase reporter assays for chemical-induced genotoxicity and oxidative damage. Applied Microbiology and Biotechnology 101, 659–671.

37. Shinohara, A., Ogawa, H., and Ogawa, T. (1992). RAD51 protein involved in repair and recombination in Saccharomyces cerevisiae is a RecA-like protein. Cell 69, 457–470.

38. Gibert, I., Calero, S., and Barbé, J. (1990). Measurement of in vivo expression of nrdA and nrdB genes of Escherichia coli by using IacZ gene fusions. Molecular and General Genetics 220, 400–408.

39. Maslowska, K.H., Makiela-Dzbenska, K., and Fijalkowska, I.J. (2019). The SOS system: A complex and tightly regulated response to DNA damage. Environmental and Molecular Mutagenesis 60, 368–384.

40. Sousa, C., De Lorenzo, V., and Cebolla, A. (1997). Modulation of gene expression through chromosomal positioning in Escherichia coli. Microbiology 143, 2071–2078.

41. Odsbu, I. Morigen, and Skarstad, K. (2009). A reduction in ribonucleotide reductase activity slows down the chromosome replication fork but does not change its localization. PLoS ONE 4, e7617.

42. Mahata, T., Molshanski-Mor, S., Goren, M.G., Jana, B., Kohen-Manor, M., Yosef, I., Avram, O., Pupko, T., Salomon, D., and Qimron, U. (2021). A phage mechanism for selective nicking of dUMP-containing DNA. Proceedings of the National Academy of Sciences 118, e2026354118.

43. Jinadasa, R.N., Bloom, S.E., Weiss, R.S., and Duhamel, G.E. (2011). Cytolethal distending toxin: a conserved bacterial genotoxin that blocks cell cycle progression, leading to apoptosis of a broad range of mammalian cell lineages. Microbiology 157, 1851–1875.

44. Naito, T., Kusano, K., and Kobayashi, I. (1995). Selfish behavior of restriction-modification systems. Science 267, 897–899.

45. Kobayashi, I. (2001). Behavior of restriction-modification systems as selfish mobile elements and their impact on genome evolution. Nucleic Acids Research 29, 3742–3756.

